# A large-scale comparison shows that genetic changes causing antibiotic resistance in experimentally evolved *Pseudomonas aeruginosa* predict those in naturally evolved bacteria

**DOI:** 10.1101/674531

**Authors:** Samuel J. T. Wardell, Attika Rehman, Lois W. Martin, Craig Winstanley, Wayne M. Patrick, Iain L. Lamont

## Abstract

*Pseudomonas aeruginosa* is an opportunistic pathogen that causes a wide range of acute and chronic infections. An increasing number of isolates have acquired mutations that make them antibiotic resistant, making treatment more difficult. To identify resistance-associated mutations we experimentally evolved the antibiotic sensitive strain *P. aeruginosa* PAO1 to become resistant to three widely used anti-pseudomonal antibiotics, ciprofloxacin, meropenem and tobramycin. Mutants were able to tolerate up to 2048-fold higher concentrations of antibiotic than strain PAO1. Genome sequences were determined for thirteen mutants for each antibiotic. Each mutant had between 2 and 8 mutations. There were at least 8 genes mutated in more than one mutant per antibiotic, demonstrating the complexity of the genetic basis of resistance. Additionally, large deletions of up to 479kb arose in multiple meropenem resistant mutants. For all three antibiotics mutations arose in genes known to be associated with resistance, but also in genes not previously associated with resistance. To determine the clinical relevance of mutations uncovered in experimentally-evolved mutants we analysed the corresponding genes in 457 isolates of *P. aeruginosa* from patients with cystic fibrosis or bronchiectasis as well as 172 isolates from the general environment. Many of the genes identified through experimental evolution had changes predicted to be function-altering in clinical isolates but not in isolates from the general environment, showing that mutated genes in experimentally evolved bacteria can predict those that undergo mutation during infection. These findings expand understanding of the genetic basis of antibiotic resistance in *P. aeruginosa* as well as demonstrating the validity of experimental evolution in identifying clinically-relevant resistance-associated mutations.

**Importance:** The rise in antibiotic resistant bacteria represents an impending global health crisis. As such, understanding the genetic mechanisms underpinning this resistance can be a crucial piece of the puzzle to combatting it. The importance of this research is that by experimentally evolving *P. aeruginosa* to three clinically relevant antibiotics, we have generated a catalogue of genes that can contribute to resistance *in vitro*. We show that many (but not all) of these genes are clinically relevant, by identifying variants in clinical isolates of *P. aeruginosa*. This research furthers our understanding of the genetics leading to resistance in *P. aeruginosa* and provides tangible evidence that these genes can play a role clinically, potentially leading to new druggable targets or inform therapies.

## Introduction

*Pseudomonas aeruginosa* is an opportunistic pathogen responsible for a wide range of acute and chronic infections. It is a frequent cause of hospital acquired infections and pneumonia (1–3). *P. aeruginosa* chronically infects the lungs of most adults with cystic fibrosis (CF) as well as patients with bronchiectasis and chronic obstructive pulmonary disease. Despite ongoing and regular treatment with anti-pseudomonal antibiotics, infections commonly persist resulting in reduced quality of life and premature death for these patients (4–7). The failure of antibiotic therapy to eradicate chronic *P. aeruginosa* infection from the airways is a contributing factor to development of antibiotic resistance within bacteria infecting these patients (8, 9). It arises in part because the bacteria adopt a biofilm lifestyle and also because the anatomically distinct regions within the lung lead to multiple different microenvironments. Consequently many bacteria are likely to experience sub-inhibitory concentrations of antibiotics with selection for more highly resistant mutants (10).

The basis of antibiotic resistance in *P. aeruginosa* has been investigated using genetic and biochemical approaches. Mechanisms of resistance can be classified into four groups: reduced antibiotic uptake, enhanced antibiotic efflux, reduced affinity of antibiotics to their cellular targets, and inactivation of antibiotics (11–13). Resistance mainly occurs through mutations, although genes obtained via horizontal gene transfer also confer a resistant phenotype.

The emergence of whole genome sequencing (WGS) technologies has provided new approaches to understanding the molecular mechanisms driving antibiotic resistance (14, 15). Experimental evolution of antibiotic-resistant mutants from sensitive parent strains, followed by whole genome sequencing to identify resistance-conferring mutations, is an approach that has been applied to a number of species (16–21), including *P. aeruginosa* (22–28). Studies have confirmed the involvement of genes previously proposed to be associated with resistance in *P. aeruginosa*. These include the *fusA1* gene that encodes the EFG-1 protein and is associated with aminoglycoside resistance (27, 29, 30) and the *ftsI* gene that encodes a penicillin binding protein and is associated with carbapenem resistance (30–34). This approach has high potential for identifying previously unknown antibiotic resistance-associated mutations and genes, but some key issues remain to be addressed. Experimental evolution of small numbers of mutants may overlook mutations that can contribute to resistance but do not always arise, making it more difficult to draw robust conclusions. The mutations obtained may be influenced by the selection method used – for example, continuous exposure to increasing amounts of antibiotic in broth culture may give different outcomes to intermittent antibiotic exposure, as occurs during infection in patients. Lastly and perhaps most importantly a rigorous comparison of experimentally evolved bacteria and those that have evolved naturally during infection is lacking despite the possibility of resistance mutations in experimentally evolved bacteria differing from those that arise in bacteria during infection. For example, mutations that increase resistance in the laboratory setting may not be tolerated in the complex environment of an infection. It is also of clinical importance to determine whether experimentally evolved mutants have cross resistance or increased susceptibility (*i.e.* collateral sensitivity) to other antibiotics. The overall aim of this research was to address these issues, and to determine the relevance of mutations arising during experimental evolution to the evolution of antibiotic resistance during infection.

## RESULTS

### Antibiotic resistance of experimentally evolved mutants

Highly resistant mutants of *P. aeruginosa* PAO1 were evolved in thirteen parallel experiments for each of three antibiotics, tobramycin, meropenem and ciprofloxacin. Each mutant was selected through serial passage on antibiotic gradient agar plates containing increasing amounts of antibiotic, interspersed with growth in antibiotic-free broth. Mutants were considered to have reached maximum resistance when selection failed to give rise to any further increase in resistance (typically between 6 and 8 serial passages). A control culture was passaged 6 times in the absence of antibiotic.

Figure 1 shows the MIC values for each of the 39 experimentally evolved mutants when tested against six different antibiotics belonging to three different classes. Compared to the parental strain *P. aeruginosa* PAO1 the evolved mutants had a minimum of 64-fold and a maximum of 2048-fold increase in their MIC values for the selecting antibiotic. All mutants had similarly increased resistance to a second antibiotic of the same class (levofloxacin, imipenem and gentamicin for ciprofloxacin, meropenem and tobramycin-selected mutants, respectively).

**Fig. 1.**
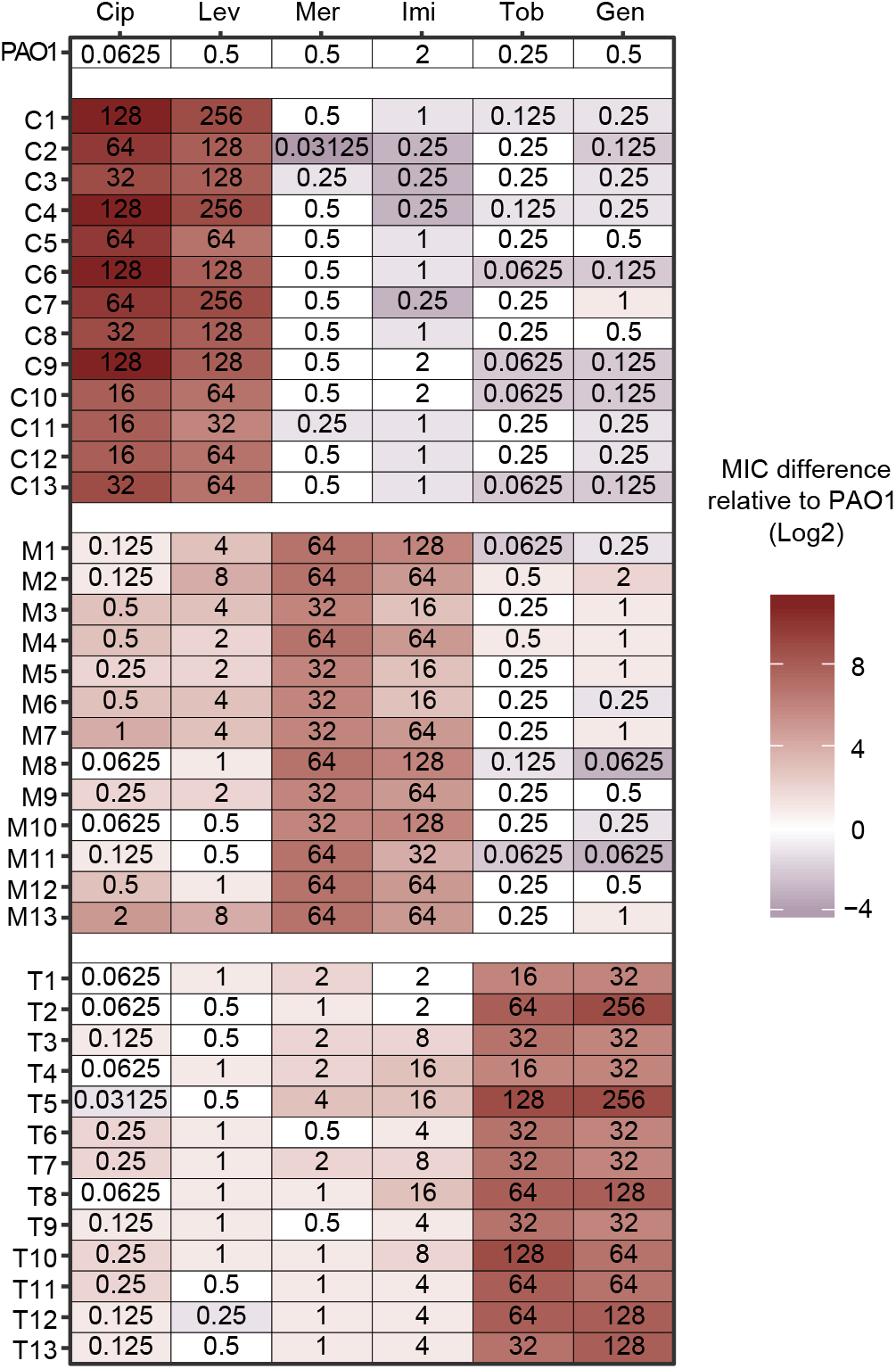
Minimum Inhibitory Concentrations of experimentally evolved antibiotic resistant mutants to fluoroquinolones, carbapenems and aminoglycosides. MIC values are shown for the parental strain *P. aeruginosa* PAO1 and for mutants selected for resistance to ciprofloxacin (C1-C13), meropenem (M1-M13) or tobramycin (T1-T13). MIC values are shown in mg/L, and coloured based on log_2_-fold change from strain PAO1, ranging from −4 (grey) to 11 (red). Clinical breakpoints as defined by EUCAST (www.eucast.org) in mg/L are: ciprofloxacin (Cip) ≥0.5, levofloxacin (Lev) ≥1, meropenem (Mer) ≥8, imipenem (Imi) ≥8, tobramycin (Tob) ≥4 and gentamicin (Gen) ≥4. A control culture that underwent six serial passages in the absence of any antibiotic selection had the same MIC for all antibiotics as the parental *P. aeruginosa* PAO1 strain.

The mutants were also tested for cross-resistance to antibiotics of different classes. Most of the meropenem-selected mutants had increased resistance to fluoroquinolone antibiotics and were at or above the EUCAST clinical breakpoint for resistance in many cases (Fig. 1). Similarly, most of the tobramycin-selected mutants had increased resistance to the carbapenems tested, in particular imipenem. Conversely, most of the ciprofloxacin-selected mutants had increased (collateral) sensitivity to at least one carbapenem or aminoglycoside and some of the meropenem-selected mutants also had increased aminoglycoside sensitivity.

### Growth of antibiotic-resistant mutants

To assess the impact of antibiotic resistance on bacterial growth, we undertook growth analysis for each of the experimentally evolved mutants (Fig. 2 and Figs. S1 and S2). Growth was measured as Area Under Growth Curve (AUC) in order to capture differences in both growth rate and bacterial cell density in stationary phase. In the absence of antibiotics, mutants in all three resistance groups grew significantly less well than the parental strain *P. aeruginosa* PAO1, with tobramycin-selected mutants showing the greatest reduction in growth (median values of 1147.5, 747.5, 610.3, and 468 AUC units for strain PAO1 and for ciprofloxacin, meropenem and tobramycin-resistant mutants respectively). However, there was a large degree of heterogeneity in growth within each group, with some mutants in each group having at least twice the growth of others in the same group.

**Fig. 2.**
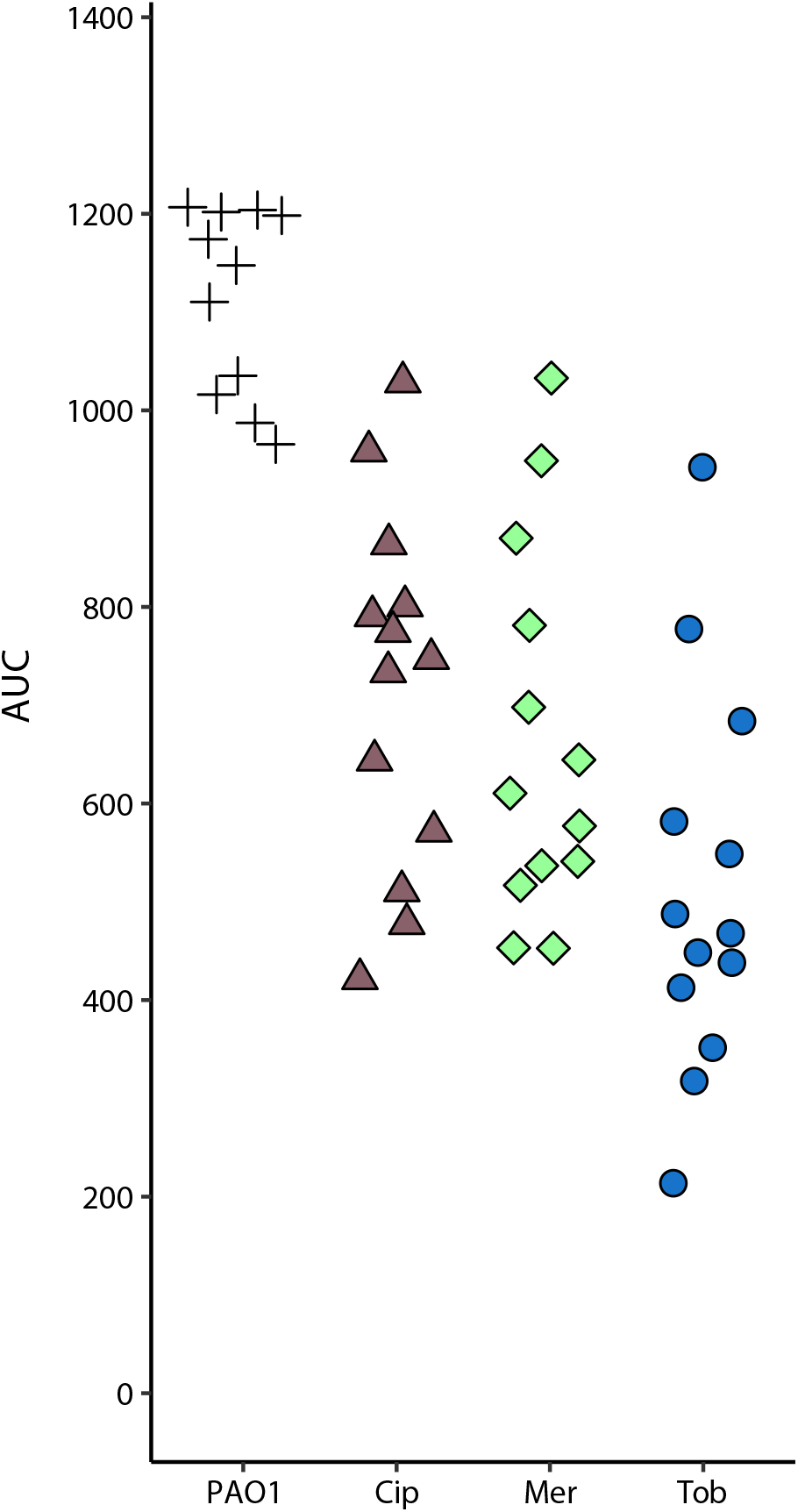
Growth analysis of experimentally evolved mutants in the absence of antibiotics. Growth of each experimentally evolved mutant was measured during 18h incubation at 37°C and Area Under Growth Curve (AUC) values determined. At least 3 biological replicates were carried out for each mutant, with mean AUC values shown. A one-way ANOVA with post-hoc Dunnett’s test was carried out on each antibiotic selection using PAO1 as a comparison (n=11). Bonferroni corrected P-values for Cip-, Mer-, and Tob-evolved mutants are 3.98×10^−6^, 3.36×10^−7^, and 2.11×10^−10^ respectively. Cip, ciprofloxacin; Mer, meropenem; Tob, tobramycin.

### Identification of antibiotic resistance associated mutations

Whole genome sequencing (WGS) and variant calling of the 39 antibiotic resistant mutants showed that each mutant contained between 2 and 8 mutations. Mutations were present across 78 genes; 27 genes were mutated in 2 or more mutants (Table 1). In addition, four intergenic mutations were present in meropenem resistant mutants and 5 of the 13 meropenem-resistant mutants contained large deletions ranging in size from 225 to 479 kb (Table S1 and Fig. S3). Putative large duplications up to 600kb were identified in 2 tobramycin and 1 meropenem resistant isolate (Fig. S4). No mutations were present in bacteria serially passaged in the absence of antibiotic selection.

**Table 1.** Genes mutated in more than one experimentally evolved antibiotic resistant mutant.

| Gene | Experimentally evolved mutants (n = 13) |
| --- | --- |
| **Ciprofloxacin resistance** |  |
| *gyrA* | 13 |
| *nfxB* | 12 |
| PA3491 | 11 |
| *parC* | 6 |
| *parE* | 3 |
| *pilT* | 3 |
| *pilF* | 3 |
| *tpiA* | 2<sup>†</sup> |
| PA5001 | 2 |
| **Meropenem resistance** |  |
| *oprD* | 11 |
| *nalC* | 6 |
| Large deletion | 5 |
| *aroB* | 3 |
| *ftsI* | 3 |
| *nalD* | 3 |
| *msrA* | 2 |
| *mexR* | 2<sup>‡</sup> |
| *phoQ* | 2 |
| **Tobramycin resistance** |  |
| *fusA1* | 13 |
| *wbpL* | 7 |
| *Δcco* | 6 |
| *amgS* | 4 |
| PA1767 | 4 |
| *mexY* | 4 |
| *rplF* | 2 |
| PA4429 | 2 |
| PA5528 | 2 |
| Putative large duplication | 2<sup>*</sup> |
| PA5001 | 2 |
| *nalC* | 2 |

<sup>†</sup>*tpiA* is also mutated in 1/13 meropenem resistant mutants.
<sup>‡</sup>*mexR* was also mutated in 1/13 ciprofloxacin resistant mutants.
<sup>\*</sup>Putative large duplications were also identified in 1/13 meropenem resistant mutants.

### Ciprofloxacin-selected mutants

Twenty-nine mutated genes were identified following WGS of the 13 ciprofloxacin evolved mutants. All of the mutants had a mutation in the *gyrA* gene and 9 of the mutants also had mutations in the *parC* or *parE* genes that encode DNA topoisomerase, known targets of ciprofloxacin (35). Twelve of the mutants also had mutations in *nfxB* that encodes an efflux pump regulator, with mutations in *nfxB* being known contributors to ciprofloxacin resistance (35). Ten of the mutants had mutations in pilin-encoding *pil* genes (Table 1, Table S1). The relationship between *pil* mutations and ciprofloxacin resistance is not clear, although very recently other researchers also reported an association between *pil* mutations and ciprofloxacin resistance (36). Eleven of the mutants had a mutation in a gene PA3491 that to the best of our knowledge has not previously been associated with antibiotic resistance.

### Meropenem-selected mutants

A total of 26 genes were mutated in the 13 meropenem-selected mutants, with 8 genes mutated in more than one mutant (Table 1, Table S1). Many of the mutations were in genes previously associated with meropenem resistance. Eleven of the mutants contained mutations in the porin-encoding *oprD* gene, with mutations in this gene representing a primary mechanism for carbapenem resistance (24, 37–39). Ten of the mutants had mutations in *nalC*, *nalD* or *mexR*, with mutations in these genes causing upregulation of the efflux pumps and being associated with β-lactam resistance (28, 40–42). Mutations in *ftsI*, that encodes the meropenem binding protein PBP3A, were present in 3 mutants consistent with the known role of such mutations in resistance (24, 43). Mutations were also present in genes not commonly associated with meropenem resistance. Three of the evolved mutants contained mutations within the *aroB* gene that encodes an enzyme dehydroquinate synthase required for synthesis of aromatic amino acids. Five of the mutants had mutations in genes encoding tRNA ligases, with a different gene being mutated in each case.

Five of the 13 mutants had large deletions ranging in size from 225kb up to 480kb. These deletions all overlapped, with the genome region between PA2022 – PA2208 being deleted in all the five mutants. A smaller deletion (1986bp) spanning PA2022-PA2024 was present in one mutant. The deleted region contains PA2023 (*galU*). Insertion mutations in *galU* lead to increase in resistance to meropenem, cephalosporin and aminoglycosides (38, 39, 44).

### Tobramycin-selected mutants

A total of 24 individual genes were mutated across the 13 tobramycin-selected mutants, with ten genes mutated in more than one mutant (Table 1, Table S1). All mutants had at least one mutation in *fusA1* that encodes elongation factor G. Mutations in *fusA1* have recently been reported to confer tobramycin resistance in laboratory-evolved mutants (29). Nine mutants also had mutations in the *wbpL* or PA5001 genes that are associated with lipopolysaccharide synthesis and ten mutants had mutations in genes directly involved in oxidative phosphorylation and generation of proton motive force (*cco*, *nuoG*, PA1549, and PA4429). Mutations affecting either of these processes increased aminoglycoside resistance in a whole-genome screen for resistance genes (45). Several other genes were mutated in a smaller number of tobramycin-selected mutants (Table S1). Many of these including PA1767, *mexY*, *amgS* and *nalC* have previously been implicated in aminoglycoside resistance (45–48).

### Frequencies of mutations in clinical isolates of *P. aeruginosa*

It is common for mutations that confer antibiotic resistance in *P. aeruginosa* to arise during infection in cystic fibrosis patients (49). We therefore addressed the question, do mutations that arose in the experimental evolution experiments reflect those that have arisen in clinical isolates of *P. aeruginosa*? To do so we analysed the genomes of 457 *P. aeruginosa* clinical isolates. For comparison we analysed the genomes of 172 *P. aeruginosa* isolated from the general environment, and which are therefore unlikely to have had antibiotic exposure. All genes that were mutated in two or more of our experimentally evolved mutants were analysed in each genome to determine whether likely resistance-associated mutations had occurred. The absence of genome sequences of ancestral strains prevented direct identification of mutations. Instead, genetic variants that are likely to alter protein function by contributing to antibiotic resistance were inferred using PROVEAN, a widely used tool for predicting the likelihood that amino acid differences affect protein function (50). The results are summarised in Table 2.

**Table 2.**
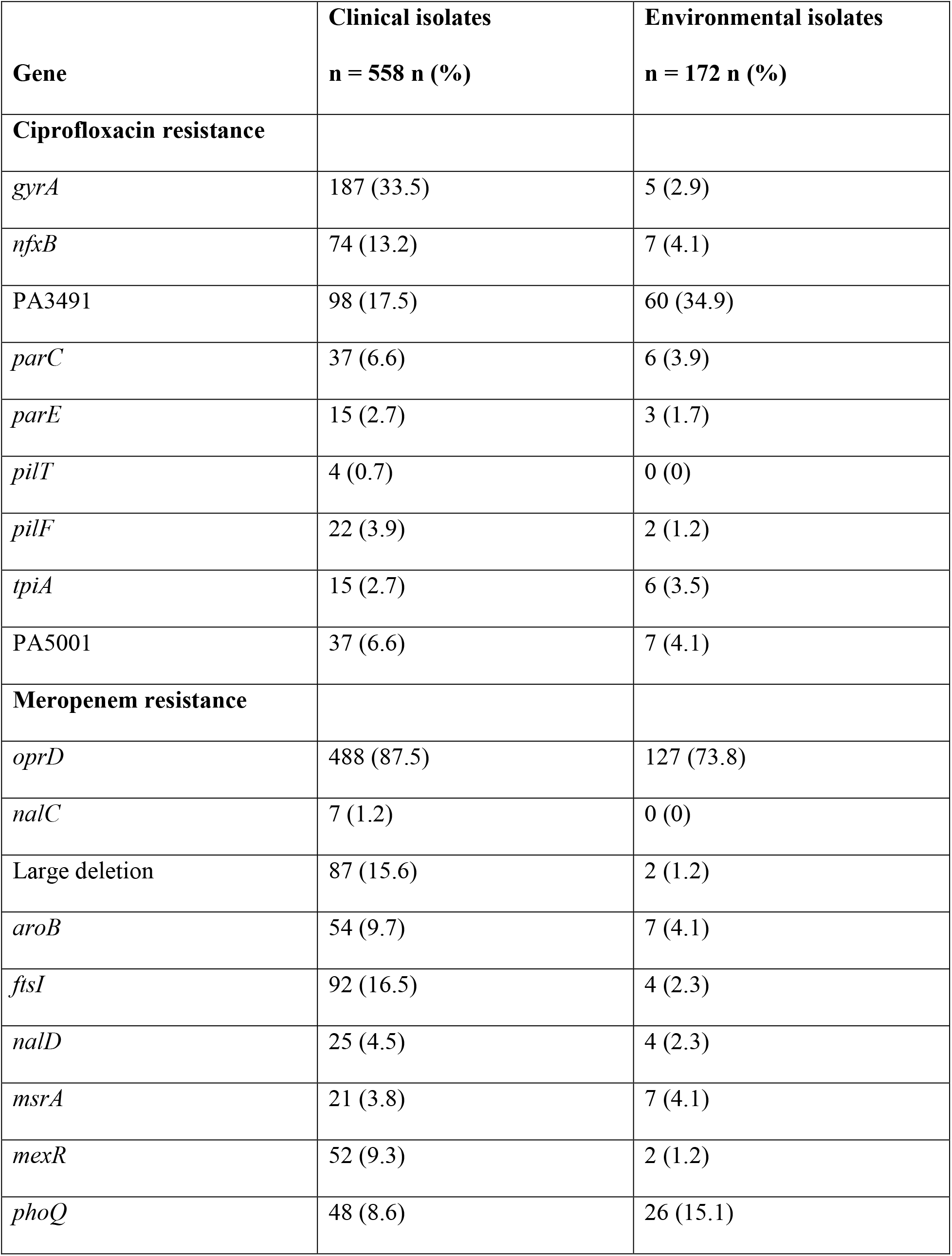
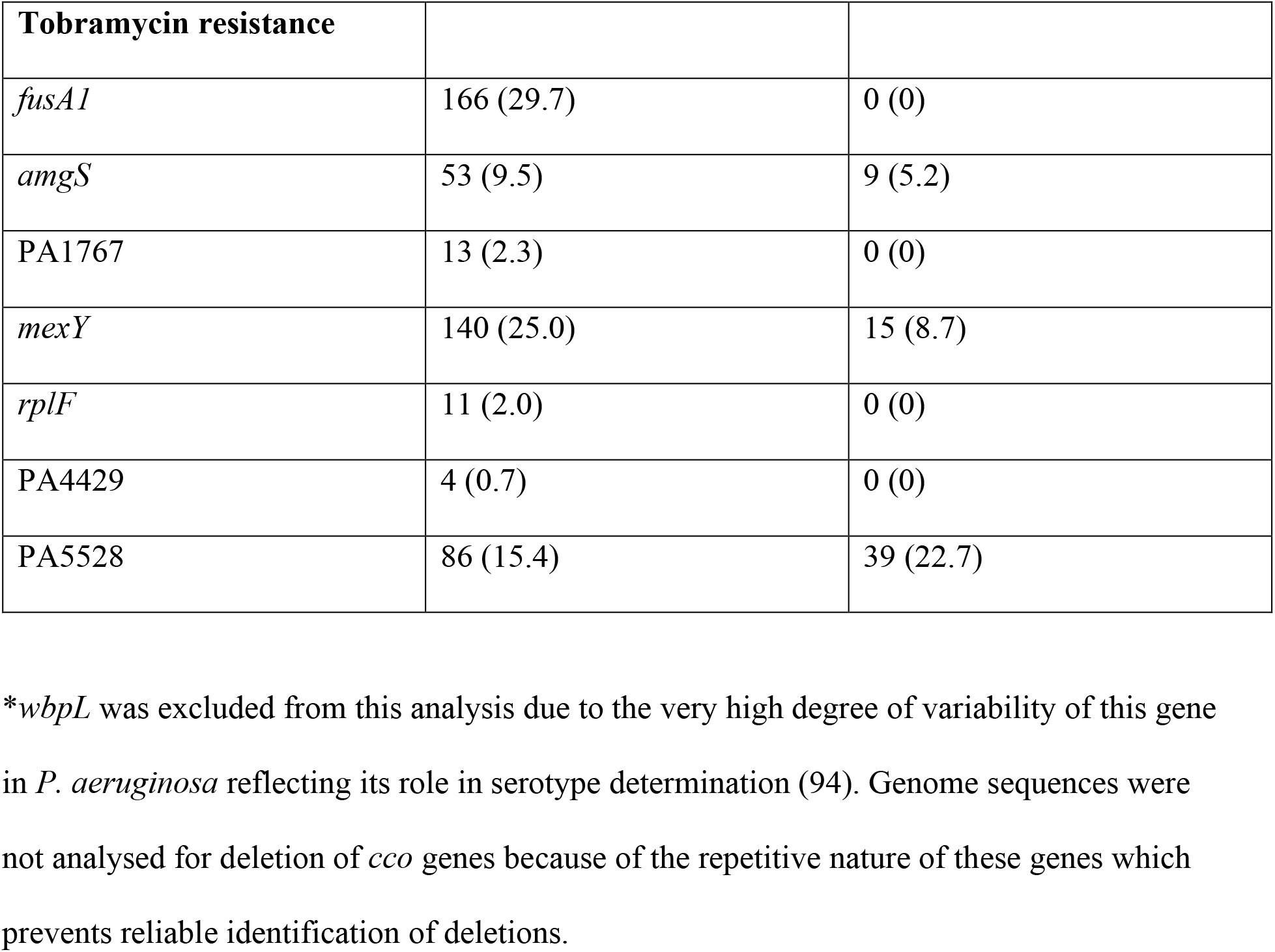
Frequencies of predicted change of function mutations in clinical and environmental isolates of *P. aeruginosa*, for genes identified through experimental evolution*.

Many of the genes that were mutated in our experimentally evolved mutants had higher frequencies of predicted function-altering variants in clinical isolates than in environmental isolates. For example, the antibiotic target genes *gyrA* (ciprofloxacin resistance), *ftsI* (meropenem resistance) and *fusA1* (tobramycin resistance) all had predicted function-altering variants in over 15% of the clinical isolates. In many cases the variations in the clinical isolates were identical to those in experimentally evolved mutants. For example, the variant T83I in GyrA was present in 96 (51.3% of isolates containing predicted functional variants) clinical isolates. Predicted function-altering variants in genes encoding regulatory proteins that are associated with antibiotic resistance such as *nalD*, *mexR* and *amgS* were also common in the clinical isolates. Predicted differences in *mexY*, which encodes an efflux pump component known to be associated with tobramycin resistance (47), were also frequent in the clinical isolates. Furthermore, the *aroB* gene, which does not have a characterised role in antibiotic resistance, was identified through whole genome sequencing of the experimentally-evolved meropenem-resistant mutants and was predicted to have function-affecting differences in over 10% of the clinical isolates. Some genes (*pilT*, *pilF*, *nalC* and PA1767) that were mutated in our experimentally-evolved mutants had only low (<5%) frequencies of predicted function-altering differences in the clinical isolates implying that mutations in these genes may not be advantageous during infection. Overall however, predicted change-of-function variants in the genes analysed had significantly less impact on the genomes of the environmental isolates than on the genomes of clinical isolates (p = 1.8×10^−11^) (Fig. 3).

**Fig. 3.**
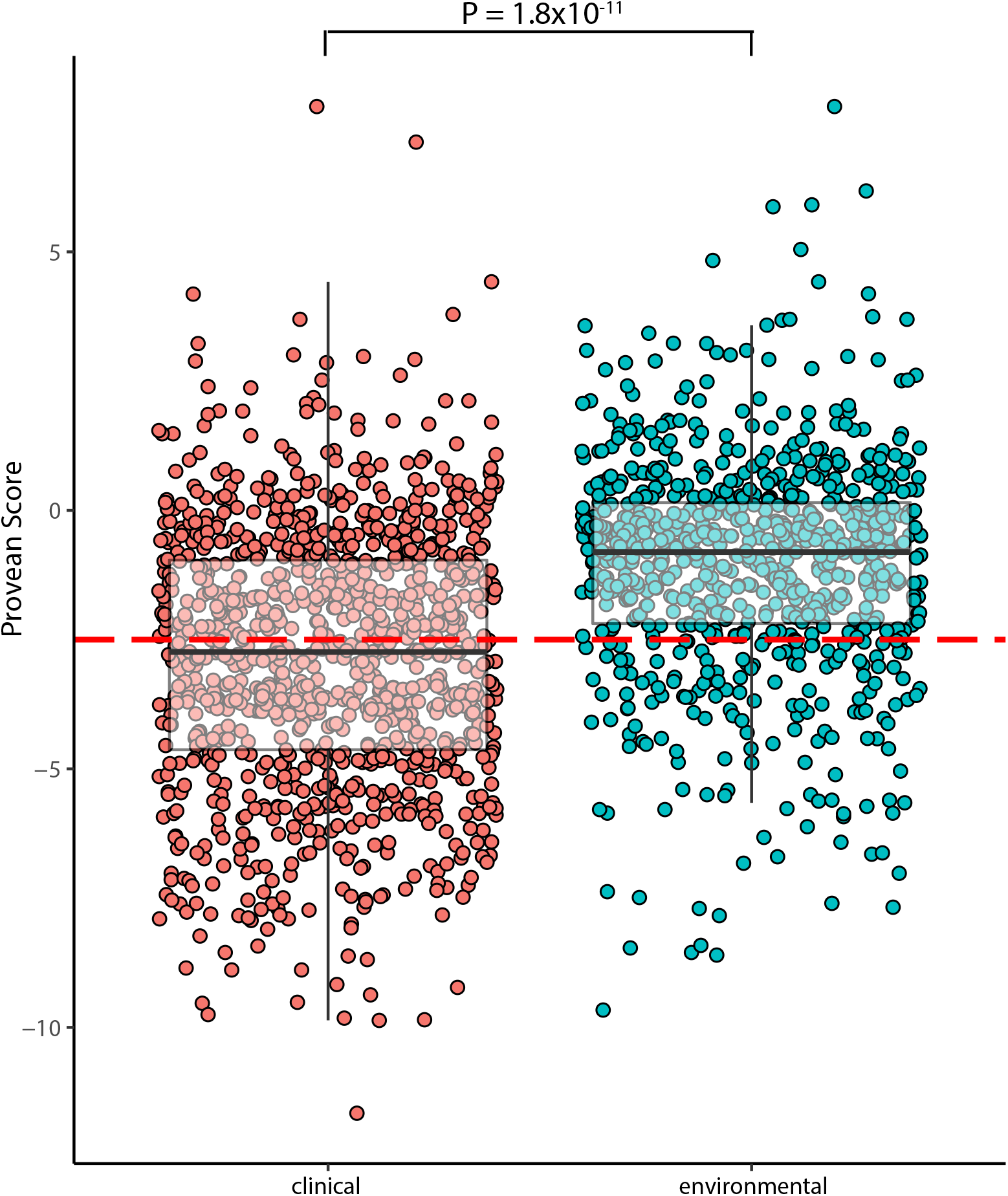
Comparison of change-of-function variants in clinical and environmental isolates. Provean values were determined for clinical and environmental (ENV) isolates for all variants in genes that were mutated in at least two experimentally evolved antibiotic resistant mutants. More negative values indicate variants that are more likely to affect function. The dashed line represents cut-off value of −2.5, with variants below this value being highly likely to affect protein function. The medians and first and third quartiles are shown in the overlaid boxplots (whiskers showing 1.5 times the interquartile range above the 75^th^ percentile and below the 25^th^ percentile). An ANOVA test using TukeyHSD correction showed that there was a significant difference in the Provean values in the clinical group compared to the ENV group (p = 1.7 × 10^−9^).

In addition to point mutations, five of the experimentally-evolved meropenem-resistant mutants had large (over 200 kb) deletions. Large deletions of comparable size were also present in 87 of the clinical isolates and only 2 of the environmental isolates (Fig. S5, Table 2).

## Discussion

In this study, we have extended the understanding of the genetics of antibiotic resistance in *P. aeruginosa*. In contrast to previous experimental evolution studies (22, 24–26, 36, 43, 51–53) we used an agar-based selection method interspersed with growth in antibiotic-free broth. This methodology, in conjunction with analysis of a large number of resistant mutants for each of three clinically-relevant antibiotics, identified a number of resistance-associated genes not found in other studies. Importantly, comparison with the genomes of *P. aeruginosa* isolated from chronically-infected patients as well as isolates from the general environment has confirmed that many mutations in experimentally-evolved mutants are clinically relevant while indicating that some appear to be restricted to the laboratory situation.

Our approach of carrying out antibiotic selection on agar plates interspersed with periods of antibiotic-free growth has some parallels with the conditions faced by *P. aeruginosa* when it colonises the lungs of cystic fibrosis patients. In both circumstances bacterial growth is on a semi-solid surface with intermittent exposure to antibiotic and so selects for mutations that are stably inherited in the absence of antibiotics. Many of the mutations identified here were in genes that were identified in other experimental evolution studies. For example, mutations altering the target site proteins GyrA and ParC as well as the efflux pump regulator NfxB are well established contributors to fluoroquinolone resistance (35). Our selection protocol and the large number of experimentally evolved mutants that we analysed also allowed us to identify some genes that were known to contribute to clinical resistance but not identified in previous experimental evolution studies. These included *parE* (ciprofloxacin resistance) (54) and *amgS* and *wbpL* (tobramycin resistance) (55).

Importantly, experimentally evolved mutants in this study also had mutations in genes not usually associated with resistance. These include PA3491 and pilin-encoding genes in ciprofloxacin-selected mutants; *aroB* and tRNA ligases in meropenem-selected mutants (39, 43); and PA1767, *cco* and *wbpL* in tobramycin-selected mutants. The occurrence of mutations in these genes in multiple independently-evolved lines (Table 1, Table S1) strongly suggests that the mutations increase antibiotic tolerance in our selection system. Entry of tobramycin into *P. aeruginosa* can be affected by changes to the cell surface or to membrane potential (45, 56) and mutations in cytochrome C oxidase or *wbpL* may result in such changes. How the other genes may affect resistance is not clear. Further research will be required to determine the how mutations in these genes increase resistance, as well as their role (if any) in resistance of clinical isolates.

Another noteworthy observation was the prevalence of large deletions (up to 8% of the genome) in mutants selected for resistance to meropenem, something that has also been observed previously (24). All the deletions overlapped, with PA2023 (*galU*) that encodes an enzyme required for LPS core synthesis being deleted in all cases. In a genome-wide screen, a mutation in *galU* increased meropenem tolerance (38, 39). Deletion of *galU* may therefore contribute to increased tolerance in the deletion-containing mutants obtained here, perhaps in conjunction with other deleted genes.

We used PROVEAN to assess the frequency of likely change-of-function mutations in isolates of *P. aeruginosa* from patients. For comparison we analysed a panel of isolates from the general environment (Table 2). A significant proportion of isolates from CF patients are antibiotic resistant, with multi-drug resistant *P. aeruginosa* found in 18% of individuals with CF (57), whereas environmental isolates are typically sensitive to antibiotics (Ramsay et al., submitted for publication).

A high proportion of clinical isolates contained likely function-altering differences in genes that are established as contributing to clinical resistance to fluoroquinolones (*gyrA* and *parE*), carbapenems (*mexR*, *nalD*) and/ or aminoglycosides (*mexY*, *amgS*) (Table2) (27, 28, 58–61). These genes contained few or no predicted function-altering differences in isolates from the general environment, emphasising their clinical relevance as well as validating our approach. Three genes *fusA1*, *ftsI* and *aroB* that have only recently been identified as affecting antibiotic resistance (*aroB* in this study) also had predicted function-altering differences in clinical isolates but few or no environmental isolates (24, 29). These findings demonstrate the utility of experimental evolution for identifying clinically-relevant genes associated with antibiotic resistance.

The *oprD* gene that is well established as being associated with carbapenem resistance had a high frequency of predicted function-altering differences in clinical isolates, as expected. It also had a high frequency of such differences in environmental isolates of *P. aeruginosa* consistent with earlier findings (37), a finding that was related to its role in environmental adaptation of *P. aeruginosa*. Nonetheless the mean PROVEAN scores for OprD were lower in the clinical isolates (−2.8 clinical, −0.8 environmental), likely reflecting the need for more severe loss of function changes to OprD in contributing to carbapenem resistance.

Conversely, the *nfxB*, *parC* and *nalC* genes that were previously shown to influence the resistance phenotype of clinical isolates (40, 54, 62) and in experimental evolution studies (Table 2) (22, 24) had only low frequencies of predicted function-altering differences in the clinical isolates in this study. This may indicate that mutations in these genes are associated with higher levels of antibiotic resistance than that of the isolates in our study. Mutations in these genes may also have a high fitness cost in the clinical environment.

A number of genes that were mutated in multiple experimentally evolved mutants are not commonly associated with antibiotic resistance. Many of these, such as *pil* genes, *tpiA* and PA1767, had only low frequencies of predicted function-changing differences in the clinical isolates. This finding suggests that mutations in these genes contribute to increased antibiotic tolerance in laboratory culture but likely do not do so during infection and further emphasises the importance of comparing experimentally-evolved mutants with clinical isolates. In contrast, mutations in genes such as *mexZ* and *gyrB* are well-characterised in clinical isolates (32, 63–66) but were not identified in our experiments or in other experimental evolution studies, indicating that experimental evolution studies alone are not necessarily sufficient to identify all resistance-associated mutations.

The experimentally evolved mutants had increased tolerance to antibiotics of the same class (Fig. 1), as found previously (25). Many of the mutants also had altered MICs for antibiotics of different classes and some of these changes could be related to the mutations that were present. For example many of the meropenem-selected mutants had mutations in *nalC* and such mutations lead to overexpression of *mexAB-oprM* (28), which may reduce susceptibility to fluoroquinolones. Conversely several of the ciprofloxacin-selected mutants had increased susceptibility to aminoglycosides and to imipenem, likely due to the presence of *nfxB* mutations (67). The occurrence of collateral changes in antibiotic sensitivity as a result of mutations selected in response to antibiotic exposure has also been observed in isolates of *P. aeruginosa* from patients (68–71). This has important clinical implications, because such outcomes during infection may affect treatment options for patients with CF or other chronic infections.

In the absence of antibiotics the majority of the experimentally-evolved mutants showed significant reductions in growth relative to the wild-type control, consistent with previous studies (Fig. 2) (23, 24, 72, 73). It is noteworthy that isolates of *P. aeruginosa* from patients are often slow growing (74) although the extent to which this phenotype is related to antibiotic resistance is not clear.

In conclusion, our study demonstrates the power of experimentally evolving multiple mutants for identifying genes that, when mutated, contribute to increased antibiotic resistance. Evolution of multiple independent mutants reveals the frequencies at which mutations arise in individual genes, which is likely to be related to the extent to which they contribute to increased resistance in our selection system. Use of an agar-based method instead of the broth-based methods of earlier studies identified genes not previously associated with resistance and showed that the spectrum of resistance mutations is influenced by the selection protocol. Crucially, analysis of resistance-associated genes in clinical isolates of *P. aeruginosa* validated the clinical relevance of genes identified only through experimental evolution approaches, while demonstrating that some genes that are mutated in laboratory-based experiments are unlikely to be relevant to resistance during infection. The extension of our approach to other antibiotics and indeed, other bacterial species, will greatly strengthen understanding of the genetic basis of antibiotic resistance in infectious bacteria.

## Materials and Methods

### *In vitro* evolution of antibiotic resistant mutants

Antibiotic gradient plates were prepared, according to methodology described by Bryson and Szybalski (75), to evolve meropenem (Penembact, Venus Remedies Limited), tobramycin (Mylan New Zealand ltd) and ciprofloxacin (Cipflox, Mylan New Zealand Ltd) resistant *P. aeruginosa* mutants. Briefly, 15mL aliquots of molten Difco™ Muller-Hinton (MH) agar were poured into a Petri dish tilted at an approximate incline of 15° to ensure a slanted slope of media. When cooled, the Petri dish was placed on a flat surface and an additional 15mL of molten MH agar, supplemented with the antibiotic treatment, was poured onto the slanted MH agar to create an antibiotic gradient across the plate. Overnight broth culture prepared from a single colony subculture of reference strain *P. aeruginosa* PAO1 was adjusted to OD_600_ = 0.01 (1.5 × 10^6^ CFU/ml) and a 2mL aliquot poured onto the antibiotic gradient plate. Excess liquid was removed by pipette following 10-minute incubation at room temperature. Inoculated plates were incubated in aerobic conditions for 24 hours at 37°C. A single colony, isolated from an area of high antibiotic concentration from the gradient plate was selected and cultured overnight in Luria broth at 37°C/200rpm. This inoculum was then adjusted as described above and used to inoculate a subsequent gradient plate containing antibiotics one doubling concentration greater than the previous plate. This technique was repeated with increasing antibiotic concentrations until it was no longer possible to identify mutants resistant to higher amounts of antibiotic maximum resistance was reached. Thirteen replicate experiments were carried out for each antibiotic, with one mutant from experiment being used for further study. As a control, strain PAO1 was passaged six times on agar without antibiotic.

### Minimum Inhibitory Concentration (MIC) testing

MIC testing was carried out in accordance with the protocol described by Wiegand and colleagues (76). Briefly, overnight broth culture of the evolved mutants and the laboratory strain PAO1 were adjusted to OD_600_= 0.01 (1.5×10^6^ CFU/mL) and 5µL aliquots spread onto MH agar plates containing doubling antibiotic concentrations. Control plates, MH agar without antibiotic supplementation, were used as a growth comparison. All MIC plates were incubated in aerobic conditions at 37°C for 24 hours. The MIC for each isolate was determined as the lowest concentration that inhibited visible growth, excluding single colonies or faint haze. Antibiotics selected for testing included each of the target antibiotics along with one other antibiotic from the same class (aminoglycosides [gentamicin], carbapenems [imipenem] and fluoroquinolones [levofloxacin]). The EUCAST guidelines (www.eucast.org) were used to interpret antimicrobial susceptibility patterns.

### Growth analysis

Overnight broth cultures of the antibiotic resistant mutants and laboratory strain PAO1 were adjusted to OD_600_= 0.01 (1.5×10^6^ CFU/mL) and 200µL aliquots were dispensed into JETbiofil® 96-well tissue culture plates. The microtiter plates were incubated in a BMG FLUOstar omega microplate reader at 37°C/200 rpm for 18 hours. Optical density (OD_600nm_) was recorded every 30 minutes to measure the growth of the isolates. Area under curve (AUC) was used to give a measure of growth that included lag phase, rate of growth during log phase, and final cell density. Growth dynamics were calculated by using the R package GrowthCurver version 0.2.1 (77). Logistic area under curve (AUC) was used as the metric for quantifying growth.

### Clinical and environmental isolates of *P. aeruginosa*

A cohort of 558 clinical and 172 environmental genome assemblies of *P. aeruginosa* isolates from multiple countries were used in this study (Table S2) (78, 79). Clinical isolates were from patients with cystic fibrosis, bronchiectasis or chronic obstructive pulmonary disease. Six isolates from patients with cystic fibrosis were unique to this study.

### Statistical analysis

All statistical analyses were carried out using R version 3.5.0 (80). Post-hoc Dunnett tests were done using R package multcomp version 1.4-8 (81). Plots were created using R package ggplot2 (82).

### Whole genome sequencing

Genome assemblies of clinical and environmental isolates have been described previously (78, 79) (Table S2). For experimentally evolved mutants and clinical isolates sequenced in this study, genomic DNA was extracted from overnight cultures of endpoint resistant mutants using the MoBio UltraClean® Microbial DNA isolation kit. Library preparation and sequencing was carried out by New Zealand Genomics Limited using Illumina HiSeq2000 and Illumina MiSeq.

### Analysis of genomic data and mutation detection

Raw sequencing reads were examined for quality pre- and post-trimming using FastQC version 0.11.5. Trimming of raw reads was done using Trimmomatic version 0.36 (83). Draft genomes of clinical isolates sequenced in this study were assembled using SPAdes version 3.12.0 (84). The genome sequence of the *P. aeruginosa* PAO1 used in this study (PAO1-Otago) was assembled using a combination of in-house code and GDtools part of the BreSeq package (85), with mapping to the *P. aeruginosa* PAO1 genome sequence at https://www.pseudomonas.com (86). A full list of differences between the genomes is shown in Table S3. Mutations in the experimentally evolved resistant isolates were identified through comparison to PAO1-Otago using BreSeq version 0.30.0 (85).

### Prediction of duplications

For prediction of duplicated regions within the experimentally evolved mutants, CNOGpro (87) was used. Hits files were generated from bam files output in the breseq analysis and parsed into CNOGpro, GC content was normalized, and 1000 bootstrap replicates were performed using quartiles of 0.025 and 0.975. HMM correction was performed allowing for a maximum of 3 possible states, allowing for an error rate of 0.01. Tables were output containing predicted copy number for each gene and intergenic region. Regions were called as putative duplications when there was a sustained increase in predicted copy number across more than 5 genes. Known multi-copy operons were removed after comparison to the wild-type *P. aeruginosa* PAO1-Otago coverage.

### Mutant comparisons to clinical and environmental *P. aeruginosa* isolates

Protein sequences of genes found to be mutated within this study were obtained from https://www.pseudomonas.com (86). These were used as query sequences for a tblastn search against compiled databases containing either clinical, or environmental *P. aeruginosa* isolates (Table S2). Blast outputs were aligned to the reference PAO1 sequence using Clustal-Omega version 1.2.4 (88). Polymorphic sites were called using a modified version of snp-sites version 2.3.2 (89). Non-synonymous polymorphic sites (variants) were extracted from vcf files. The effects of variants were predicted using Provean version 1.1 (50) compiled using BLAST version 2.2.28+ (90), psiBLAST version 2.2.28+ (91), and cd-hit version 4.7 (92), and the NCBI non-redundant database (version retrieved 1 May 2019) (93). Variations with a score of −2.5 or below were considered as being likely to affect the biological functions of proteins (50).

### Data availability

Raw sequence reads (fastq format) of experimentally-evolved mutants are available under BioProject PRJNA542028 on the NCBI Sequence Read Archive (SRA). Accession numbers for genomes of other isolates of *P. aeruginosa* are listed in Table S2.

## Acknowledgments

This research was supported by research grants to ILL and WMP from the University of Otago and the New Zealand Health Research Council (17/372). CW gratefully acknowledges support from the Cystic Fibrosis Trust (UK). SJTW was supported by a Postgraduate Scholarship from the University of Otago and AR by a New Zealand International Doctoral Research Scholarship. We are very grateful to Roger Levesque and co-workers for making genome assemblies available prior to publication. Additionally, we are grateful to all members of the International Pseudomonas Consortium Database (IPCD), who donated strains utilized in this study. We are grateful to Kay Ramsay for her comments on an earlier version of this manuscript.

---

**Supplementary Fig. S1. Growth of experimentally evolved antibiotic resistant mutants.** Bacteria were grown in L-broth and A600 measured at 30 minute intervals. Means of 3 biological replicates (5 for ciprofloxacin resistant mutants) are shown +/− standard error. **(A**) Ciprofloxacin resistant mutants. (**B)** Meropenem resistant mutants. (**C**) Tobramycin resistant mutants.

**Supplementary Fig. S2. Analysis of growth of experimentally evolved mutants.** Area under growth curve (AUC) was calculated for the mean of each of the biological replicates for each mutant (Fig. S1). Panels A, B, and C show the ciprofloxacin, meropenem, and tobramycin resistant mutants respectively. Statistical analysis was carried out using a one-way ANOVA with post-hoc Dunnett’s test using untreated PAO1 as a comparison. Bonferroni corrected P-values are indicated by * (<0.05) ** (<0.01), and *** (<0.001), with NS indicating no significant difference from strain PAO1.

**Supplementary Fig. S3. BRIG comparison of experimentally evolved meropenem resistant mutants containing large deletions.** Genome assemblies of experimentally evolved meropenem resistant mutants were compared with the parental *P. aeruginosa* PAO1 genome using BRIG. Blue rings, genome alignment of meropenem resistant mutants containing large deletions (indicated by hatching). Black rings, mutants without deletions.

**Supplementary Fig. S4. Sequencing coverage of experimentally evolved mutants containing putative duplications of large regions in the genome.** CNOGpro, examining the read depth across the *P. aeruginosa* PAO1 genome, was used. Putative duplications are indicated by increased read coverage, and deletions by absence of sequence reads. (**A**) *P. aeruginosa* PAO1 used in this study mapped to refseq *P. aeruginosa* PAO1 (NC_002516.2). Increased read coverage indicates regions which are either duplicated in our reference strain or are present in more than one copy in the genome. (**B**) Read coverage across tobramycin resistant mutant T6. (**C**) Read coverage across tobramycin resistant mutant T11. (**D**) Read coverage across meropenem resistant mutant M1 that contains 2 putative duplications.

**Supplementary Fig. 5. BRIG comparison of clinical *P. aeruginosa* isolates containing large genome deletions.** Clinical and environmental isolates of *P. aeruginosa* were compared with strain PAO1 using BRIG. Isolates containing a large deletion (>20kb) between 2mb and 2.7mb on the *P. aeruginosa* PAO1 genome were identified. Twenty five of 286 clinical isolates, but none of the environmental isolates, met these criteria and are shown in the figure.

