## Supplementary Figure S3 for "A large-scale comparison shows that genetic changes causing antibiotic resistance in experimentally evolved *Pseudomonas aeruginosa* predict those in naturally evolved bacteria"

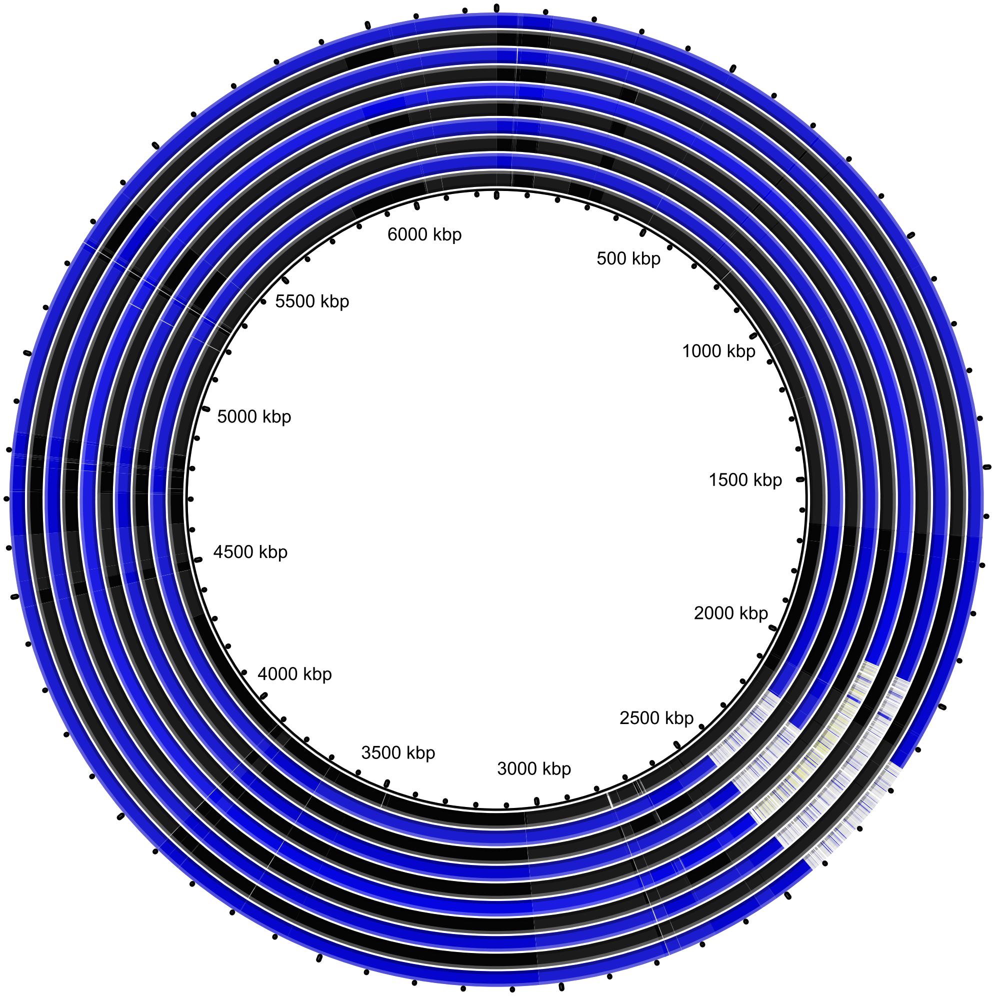


**Supplementary Fig. S3. BRIG comparison of experimentally evolved meropenem resistant mutants containing large deletions.** Genome assemblies of experimentally evolved meropenem resistant mutants were compared with the parental *P. aeruginosa* PAO1 genome using BRIG. Blue rings, genome alignment of meropenem resistant mutants containing large deletions (indicated by hatching). Black rings, mutants without deletions.
