## Supplementary Figure S4 for "A large-scale comparison shows that genetic changes causing antibiotic resistance in experimentally evolved *Pseudomonas aeruginosa* predict those in naturally evolved bacteria"

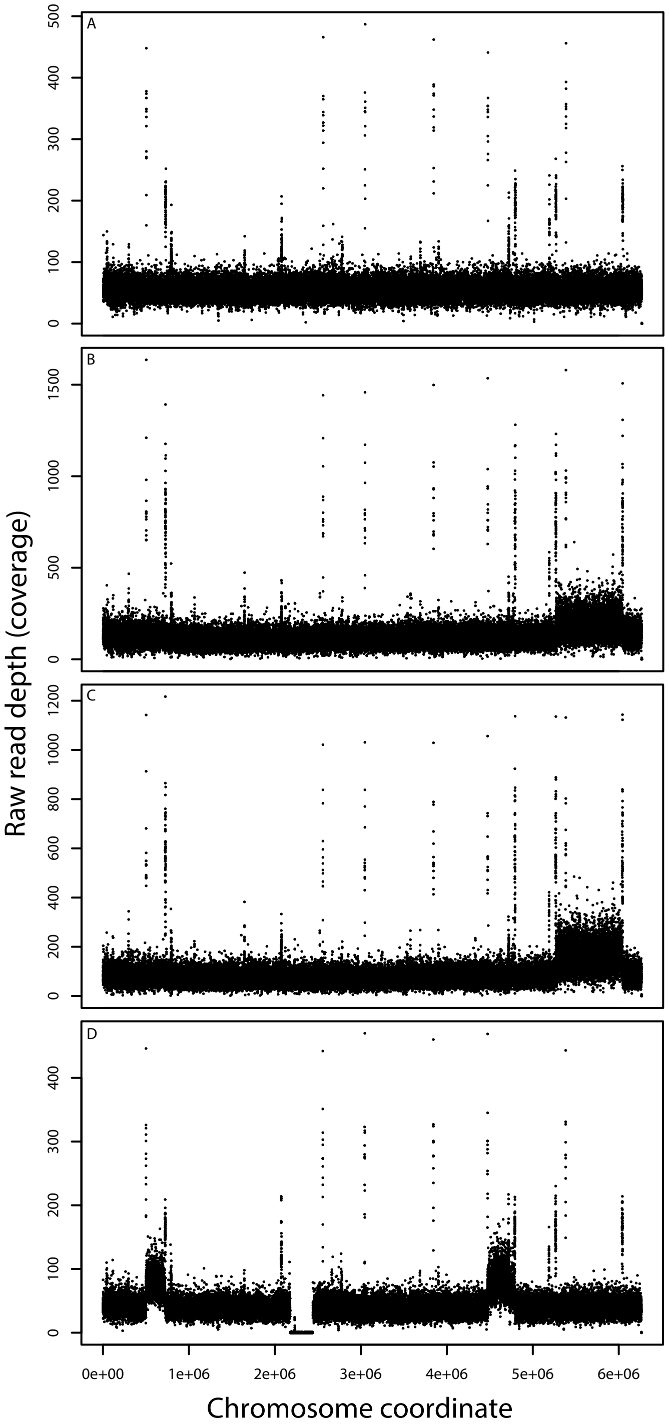


**Supplementary Fig. S4. Sequencing coverage of experimentally evolved mutants containing putative duplications of large regions in the genome.** CNOGpro, examining the read depth across the *P. aeruginosa* PAO1 genome, was used. Putative duplications are indicated by increased read coverage, and deletions by absence of sequence reads. (A) *P. aeruginosa* PAO1 used in this study mapped to refseq *P. aeruginosa* PAO1 (NC_002516.2). Increased read coverage indicates regions which are either duplicated in our reference strain or are present in more than one copy in the genome. (B) Read coverage across tobramycin resistant mutant T6. (C) Read coverage across tobramycin resistant mutant T11. (D) Read coverage across meropenem resistant mutant M1 that contains 2 putative duplications.
