## Supplementary Figure S5 for "A large-scale comparison shows that genetic changes causing antibiotic resistance in experimentally evolved *Pseudomonas aeruginosa* predict those in naturally evolved bacteria"

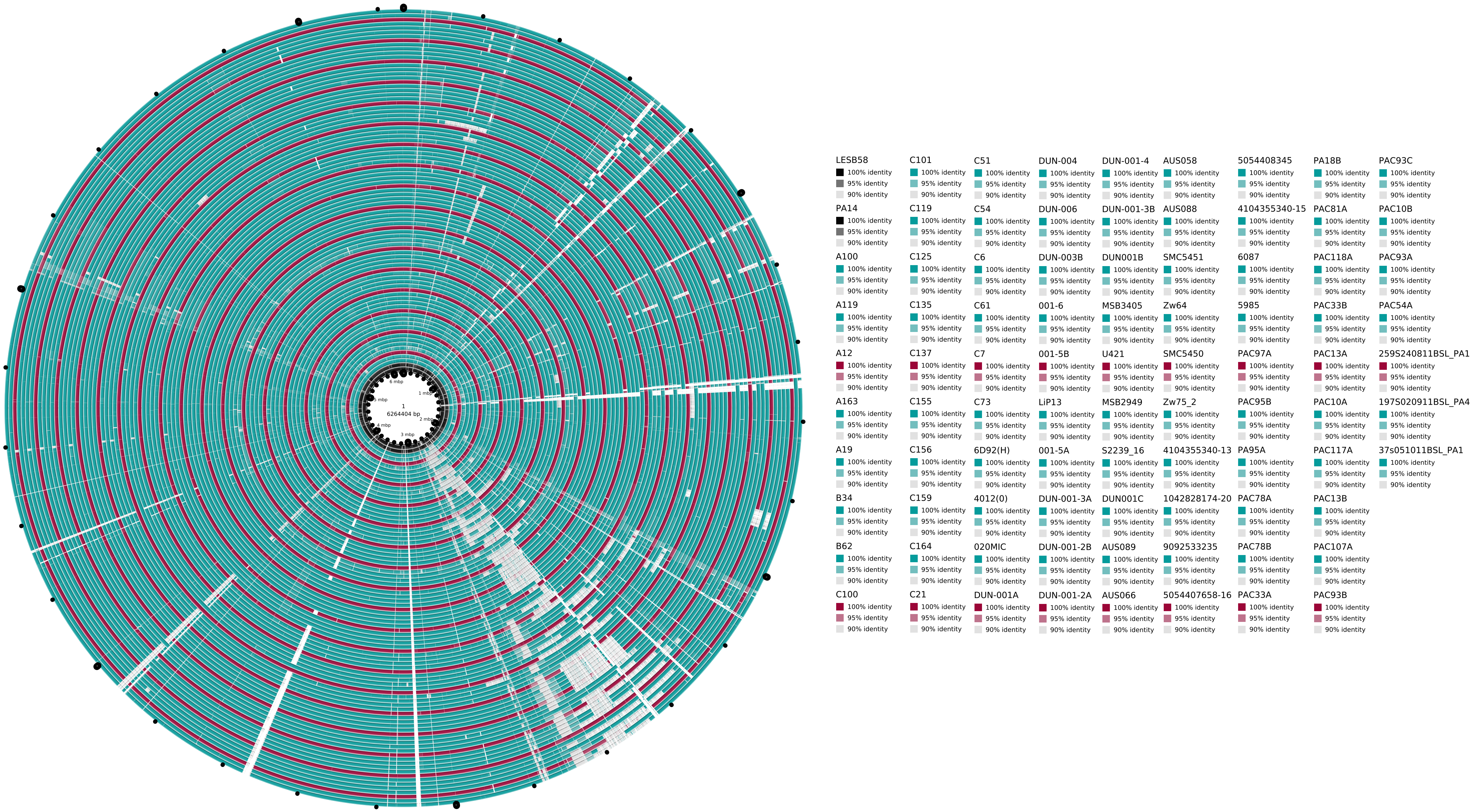


**Supplementary Fig. 5. BRIG comparison of clinical P. aeruginosa isolates containing large genome deletions.** Clinical and environmental isolates of *P. aeruginosa* were compared with strain PAO1 using BRIG. Isolates containing a large deletion (>20kb) between 2mb and 2.7mb on the *P. aeruginosa* PAO1 genome were identified. Twenty five of 286 clinical isolates, but none of the environmental isolates, met these criteria and are shown in the figure.
