## Supplementary figures and images for "A large-scale comparison shows that genetic changes causing antibiotic resistance in experimentally evolved *Pseudomonas aeruginosa* predict those in naturally evolved bacteria"

### Supplementary Figure S1

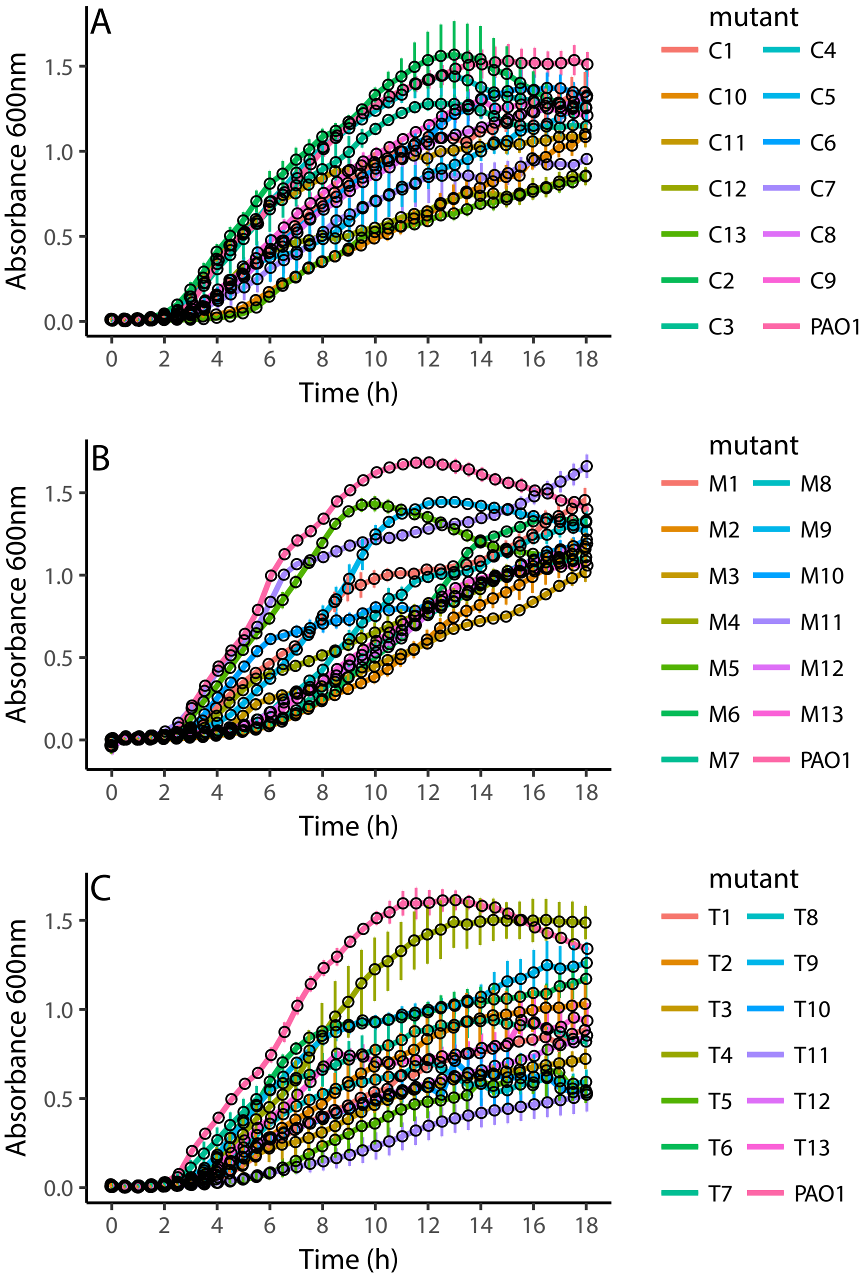

### Supplementary Figure S2

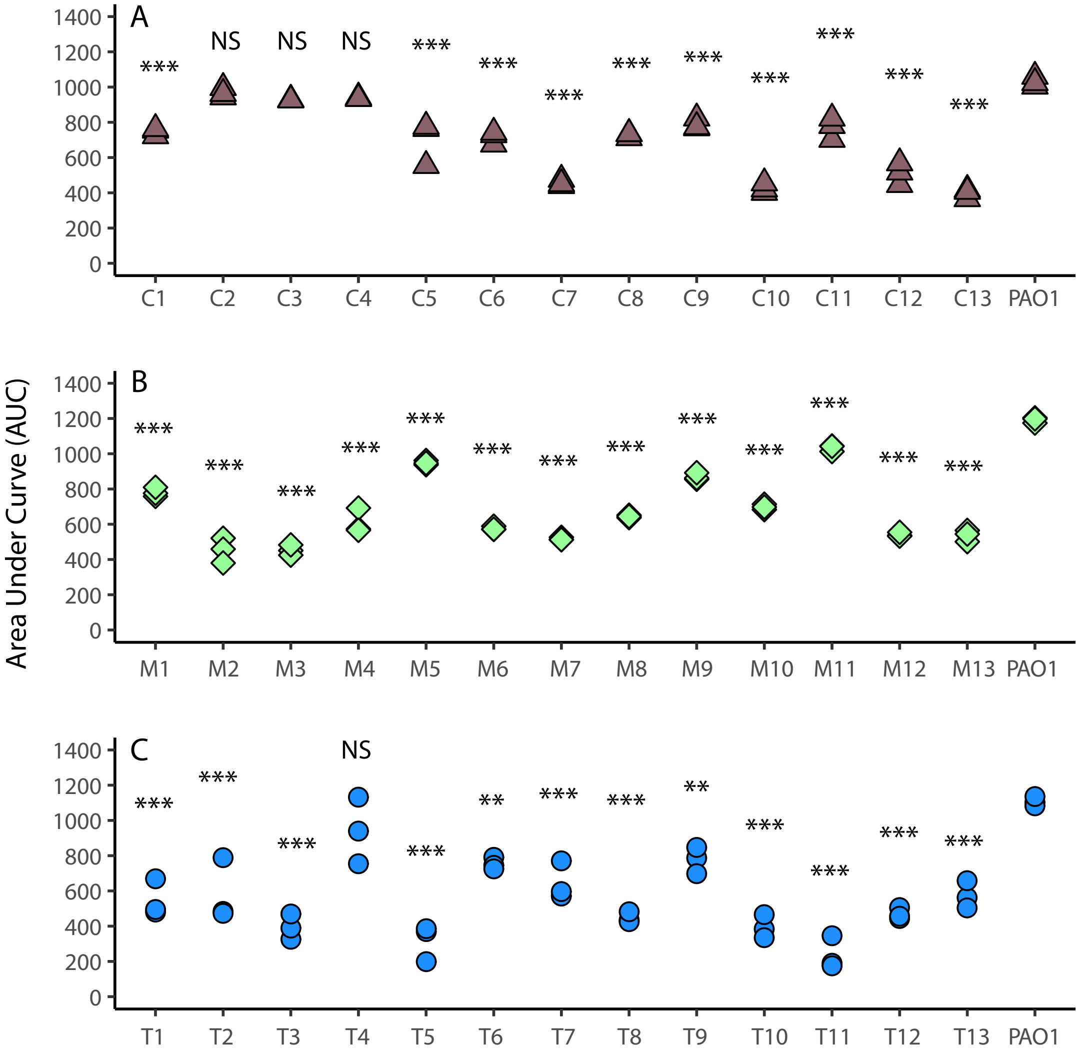
